# Zebrafish Otolith Biomineralization Requires Polyketide Synthase

**DOI:** 10.1101/396630

**Authors:** Kevin D. Thiessen, Steven J. Grzegorski, Lisa Higuchi, Jordan A. Shavit, Kenneth L. Kramer

## Abstract

Deflecting biomineralized crystals attached to vestibular hair cells are necessary for maintaining balance. Zebrafish (*Danio rerio*) are useful organisms to study these biomineralized crystals called otoliths, as many required genes are homologous to human otoconial development. We sought to identify and characterize the causative gene in a trio of mutants, *no content (nco)* and *corkscrew (csr)*, and *vanished (vns)*, which fail to develop otoliths during early ear development. We show that *nco, csr*, and *vns* have potentially deleterious mutations in polyketide synthase (*pks1*), a multi-modular protein that has been previously implicated in biomineralization events in chordates and echinoderms. We found that Otoconin-90 (Oc90) expression within the otocyst is normal in *nco* and *csr*; therefore, it is not sufficient for otolith biomineralization in zebrafish. Similarly, normal localization of Otogelin, a protein required for otolith tethering in the otolithic membrane, is not sufficient for Oc90 attachment. Furthermore, eNOS signaling and Endothelin-1 signaling were the most up- and down-regulated pathways during otolith agenesis in *nco*, respectively. Our results demonstrate distinct processes for otolith nucleation and biomineralization in vertebrates and will be a starting point for models that are independent of Oc90-mediated seeding. This study will serve as a basis for investigating the role of eNOS signaling and Endothelin-1 signaling during otolith formation.

## 1. Introduction

Otoconia and otoliths act as a mass load that increase the sensitivity of mechanosensory hair cells to the effects of gravity and linear acceleration in mammals and fish, respectively. While the morphology of otoconia (“ear particles”) and otoliths (“ear stones”) differ, the initial formation of bio-crystals rely on many homologous proteins [1]. Zebrafish otoliths are primarily composed of calcium carbonate in the form of aragonite, which accounts for ~99% of the total otolithic mass [2, 3]. The center of the otolith contains a proteinaceous core that acts as a site for otolith nucleation and biomineralization. This matrix lays the foundation for further otolith growth, which is mediated by daily deposition of additional otoconins and calcium carbonate molecules [2].

In zebrafish, otolith nucleation occurs when the otolith precursor particles or OPPs bind to the tips of the immotile kinocilia of tether cells within the otic vesicle [4, 5]. Subsequent studies have demonstrated that the critical period of otolith seeding and nucleation starts at 18-18.5 hpf and ceases by 24 hpf. [1, 4, 6-8]. In mammalian inner ear development, Otoconin-90 (Oc90; the major protein component of otoconia) is necessary for otoconial seeding and nucleation [9-11]. Oc90 can bind Otolin-1 to establish a protein-rich matrix that serves as a scaffold for subsequent deposition of calcium carbonate [12, 13]. While it is not the major protein component in zebrafish otoliths, Oc90 plays an important role in otolith seeding and early development as *oc*90-morphants do not develop otoliths [1, 14, 15]. While additional gene mutations have been identified that lead to otolith agenesis [16-20], the genes responsible for several zebrafish otolith mutants have been undetermined.

In this study, we sought to identify and characterize the causative gene in a trio of zebrafish mutants, *no content* (*nco*) and *corkscrew* (*csr*), and *vanished* (*vns*), which fail to develop otoliths during early inner ear development. We provide genetic evidence that the causative gene is polyketide synthase (*pks1;* currently *wu:fc01d11*), a candidate gene that was recently identified as a key factor of otolith biomineralization in Japanese medaka (*Oryzias latipes*) [21].

## 2. Materials and Methods

All zebrafish were maintained in a temperature-controlled (28.5°C) and light-controlled (14h on/10h off) room per standardized conditions. *nco* strain (jj149) was generated by an ENU screen and obtained from ZIRC (Eugene, OR, USA). csr was a spontaneous mutant in a *bre*-KO2/*ntl*-GFP line. *vns* was a spontaneous mutant in a AB/TL background. All protocols were approved by Creighton University and the University of Michigan Animal Care and Use Committees.

Mutant *nco* embryos and wild-type (WT) clutchmates were collected during the critical period of otolith nucleation and seeding (24 hours post fertilization, hpf) and submitted for RNA sequencing. Analysis was completed using MMAPPR (Mutation Mapping Analysis Pipeline for Pooled RNA-seq) [22]. Whole genome sequencing of *csr* was performed and analyzed using MegaMapper [23]. All sequencing was conducted at the University of Nebraska Medical Center Genomics Core Facility. Accession numbers for *nco* RNA-seq and *csr* genome sequencing will be provided during review.

WT mRNA and *pks1*^L905P^ were synthesized using mMessage Machine from a clone provided by Dr. Hiroyuki Takeda (University of Tokyo), cleaned on an RNeasy column, and subsequently injected into single-cell *csr* and *nco* embryos. Naked plasmid was injected into *vns* embryos and overall penetrance of otolith formation was determined. Site-directed mutagenesis was used to generate the mutant clone containing the causative mutation in *csr* (*pks1*^L905P^ in Japanese medaka; *pks1*^A911P^ in zebrafish). Primers used for site-directed mutagenesis were:

*pks1_L905P_Forward*: *5’-GATATGGCGTGATGTCCGGTGACAGGTTGAAGATC-3’*

*pks1_L905P_Reverse*: *5’-ATCTTCAACCTGTCACCGGACATCACGCCATATC-3’*

Pathway analysis of nco was performed using Ingenuity Pathway Analysis (IPA, http://www.ingenuity.com) [24]. The Ensembl Gene IDs for differentially expressed genes were uploaded to IPA. Cut-off for gene expression analysis was set at 0.75 RPKM. The calculated z-score indicates a pathway with genes exhibiting increased mRNA levels (positive) or decreased mRNA levels (negative). No change in mRNA levels results in a z-score of zero.

*csr, nco, and vns* samples were PCR-amplified and submitted for Sanger sequencing using the following primers:

*nco_Forward: 5’-GGGAGGATGCTTGTTGTTGG-3’*

*nco_Reverse: 5’-GTGGCCCAGAATAGGATCCA-3’*

*csr_Forward: 5’-AAGACGGGGACATGACTCAG-3’*

*csr_Reverse: 5’-TTCAACAAACAGTGCTCCGG-3’*

*vns_Forward: 5’-GCCATCATTGGAATTGGATG-3’*

*vns_Reverse: 5-GGTGTTCCAGTCCCATGAGC-3’*

*csr* and *nco* embryos were collected during key stages in ear development, fixed with hydrogel and washed in CHAPS-based CLARITY-clearing solution [25]. Embryos were decalcified with EDTA before blocking, incubating in primary and secondary antibodies diluted in PBS-Triton (0.1%), and imaging by confocal microscopy. Affinity-purified rabbit polyclonal antibodies were generated to Otogelin (CGNRVDGPSASKG; 1:1000) or Oc90 (CNTQSDTVDRKPTQSKPQ; 1:1000) by conventional methods and directly labelled before immunofluorescence. Other antibodies used were Keratan Sulfate (MZ15; 1:2000; DSHB), Hair Cell Specific-1 (HCS-1; 1:500; DSHB), and acetylated-tubulin (1:500; Sigma T6793). Phalloidin (ThermoFisher A12379) was used at a concentration of 1:500.

Mitotracker Red (ThermoFisher #M22425) was resuspended in DMSO (0.25 mM) and diluted to 200 nM in E3 embryo medium. Embryos were then incubated for 20 minutes before removing Mitotracker solution and replacing with fresh E3 embryo medium. Samples were allowed to stabilize for 30 minutes before imaging at 21 hpf. Embryos were phenotyped at 27 hpf.

To test the effects of exogenous ions on otolith formation, embryos were kept in E3 Medium until early gastrulation. Embryos were washed, dechorionated, and transferred to 1X Basic Solution (58 mM NaCl, 0.4 mM MgSO4 and 5 mM HEPES) supplemented with 0.7 mM potassium chloride, 0.6 mM calcium nitrate or 0.6 mM calcium chloride. Embryos were then transferred to fresh 1X Basic Solution with respective supplement for the remaining development. Embryos were scored by the presence or absence of otoliths at 27 hpf.

Statistical significance was calculated using Fisher’s Exact Test, G-test for Independence, and Chi-Squared Distribution.

## 3. Results

### 3.1 csr and nco are genetically-linked

The most apparent phenotype of *csr* and *nco* mutants is that they fail to form otoliths or any observable complex calcium deposits within the inner ear (Fig. 1A-C). Furthermore, the mutant larvae are homozygous lethal as the swim bladder fails to inflate (Fig. 1A’-C’) and they are unable to feed. While it is still unknown why the swim bladder fails to inflate when otoliths are absent, it is a common phenotype in other mutants with otolith agenesis [14, 16-18, 22]. Due to this commonality within *csr* and *nco* content, we sought to determine if these phenotypes would complement each other. The results of the complementation test showed that some offspring failed to develop otoliths (n = 31/106), supporting that *nco* and *csr* likely are allelic.

**Figure 1:**
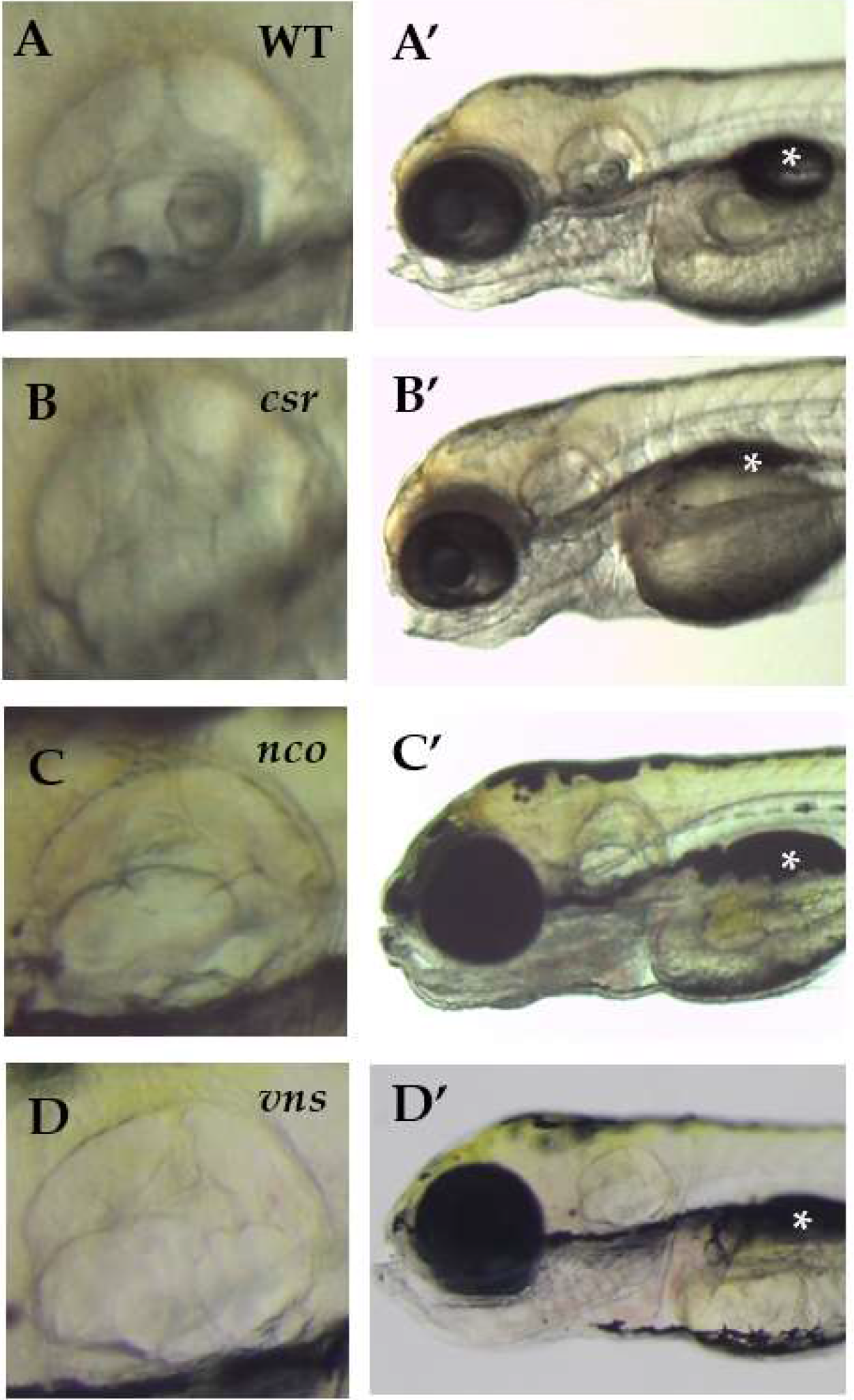
(**A-D**) The *csr, nco*, and *vns* mutant phenotypes fail to form otoliths within the inner ear. However, semicircular canal formation appears to be normal. (**A’-D’**) All mutants fail to inflate their swim bladders, which is lethal. Imaged at 5 days post fertilization (dpf). Magnification 6.3X. (*) indicates swim bladder.

### 3.2 Exogenous ions influence otolith nucleation in csr embryos; not nco or vns embryos

As an aquatic species, the environment of zebrafish can be easily controlled and adapted to assess its impact on embryonic development. Previously, small molecules have been used to block otolith development by inhibiting otolith nucleation [6]. We hypothesized that there was an error in ion homeostasis that could be affected by exogenous solutions. In water treatments supplemented with potassium chloride (n=59), we found a significant increase in *csr* embryos lacking otoliths (35.60%) compared to treatments supplemented with calcium chloride (10.94%; n=64) or calcium nitrate (20.17%; n=119)(G-test; p=0.004). Additionally, we observed no significant change in *nco* mutant embryos for water treatments supplemented with potassium chloride (17.76%; n=107), calcium chloride (16.67%; n=120) or calcium nitrate (16.9%; n=112)(G-test; p=0.975). Similarly, the penetrance of otolith formation in *vns* was not affected by exogenous salts (data not shown). Furthermore, Mitotracker was used to mark mitochondria-rich cells in *csr* and *nco* embryos. While *nco* embryos appear normal, we observed that *csr* embryos show a lack of Mitotracker localization at 21 hpf (Fig. S1). Altogether, this suggests the nature of each mutation, while likely allelic, are inherently different.

### 3.3 Potentially deleterious mutations identified in polyketide synthase for csr, nco, and vns

To positionally clone the gene responsible for *nco* and *csr*, we used complementary approaches for each strain. MMAPPR analysis of *nco*-derived RNA sequencing (Fig. 2A) [22] and MegaMapper analysis of *csr*-derived whole genome sequencing (Fig. 2B) [23] both identified a genomic region with high homology surrounding the *pks1* locus. While several other genes were in that region, a previous study on otolith biomineralization in Japanese medaka made *pks1* the likely gene candidate [21]. Potentially deleterious mutations were identified in *pks1* for *csr* (A911P) and *nco* (L681*), which were both located within a conserved acyl transferase domain (Fig. 2C). In regards to *csr* mutation, homozygosity at the locus is compatible with proper development in the AB/TL background and with otolith agenesis in the AB background (data not shown). Furthermore, a deleterious mutation in *vns* (G239R) was serendipitously found to be linked to a neighboring gene during a separate study. The deleterious point mutation was identified by Sanger sequencing of the *pks1* locus and confirmed by relatively high penetrance of otolith agenesis (95%).

**Figure 2:**
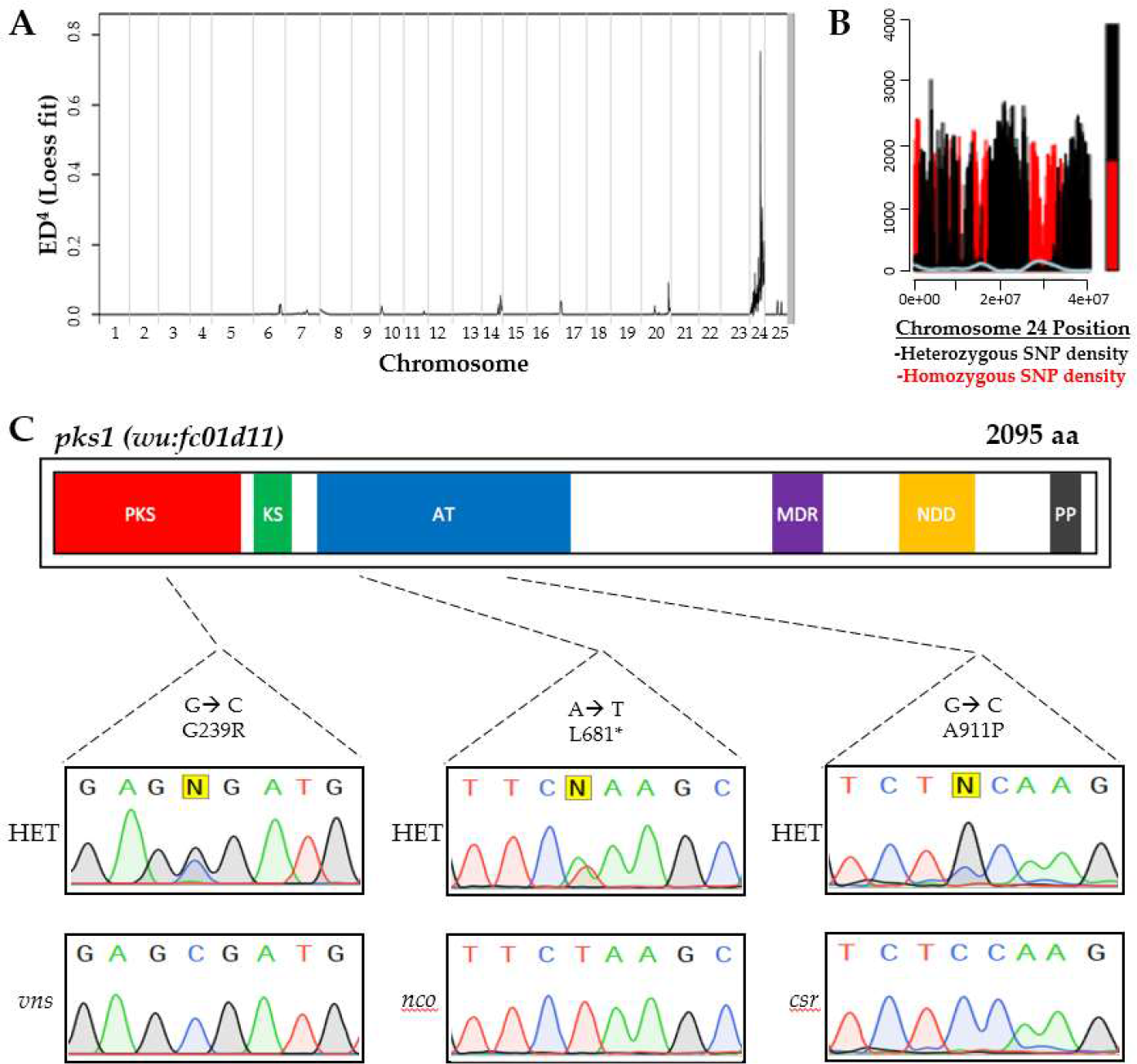
Complementary approaches for causative gene discovery. MMAPPR analysis of RNA sequencing data for *nco* (**A**) and whole genome homology mapping for *csr* (**B**) identified regions of high homology on the 24^th^ chormosome near the *pks1* locus (~33 Gb). (**C**) Deleterious mutations were identified in *pks1* for *nco* and *csr* within the acyl transferase (AT) domain and *vns* within the polyketide synthase (PKS) domain. Sanger sequencing confirmed SNPs in *csr, nco*, and *vns* mutants. Other domains include Ketoacyl Synthetase (KS), Medium Chain Reductase (MDR), NAD(P)-dependent dehydrogenase (NDD), and Phosphopanthetheine-Binding (PP).

### 3.4 Japanese medaka pks1 mRNA rescues otolith biomineralization in csr, nco, and vns

While the last common ancestor of Japanese medaka and zebrafish was estimated to be 150 million years ago [26], we sought to assess if the function of *pks1* within the inner ear is conserved. We injected Japanese medaka *pks1* mRNA or DNA into single-cell embryos of *csr, nco*, and *vns* heterozygous incrosses. Microinjection of Japanese medaka *pks1* mRNA (300 ng/μL) rescued otolith biomineralization in both *csr* (p<0.0001; χ²<0.0001; n=93) and *nco* (p=0.0032; χ²=0.0022; n=84) mutants (Fig. 3B). Additionally, microinjection of the Japanese medaka *pks1* plasmid (20 ng/uL) provided by Dr. Takeda rescued otolith biomineralization in *vns* (p<0.0001; χ²=0.0004; n=39). Using site-directed mutagenesis, we introduced the non-synonymous mutation (A911P) in *csr* to the Japanese medaka mRNA construct (L905P). We repeated injections into single-cell embryos and failed to rescue otolith biomineralization in *csr* and *nco*. WT medaka *pks1*, but not *pks1*_L905P_, rescued otolith biomineralization in *csr* and *nco* embryos (Fig. 3C).

**3:**
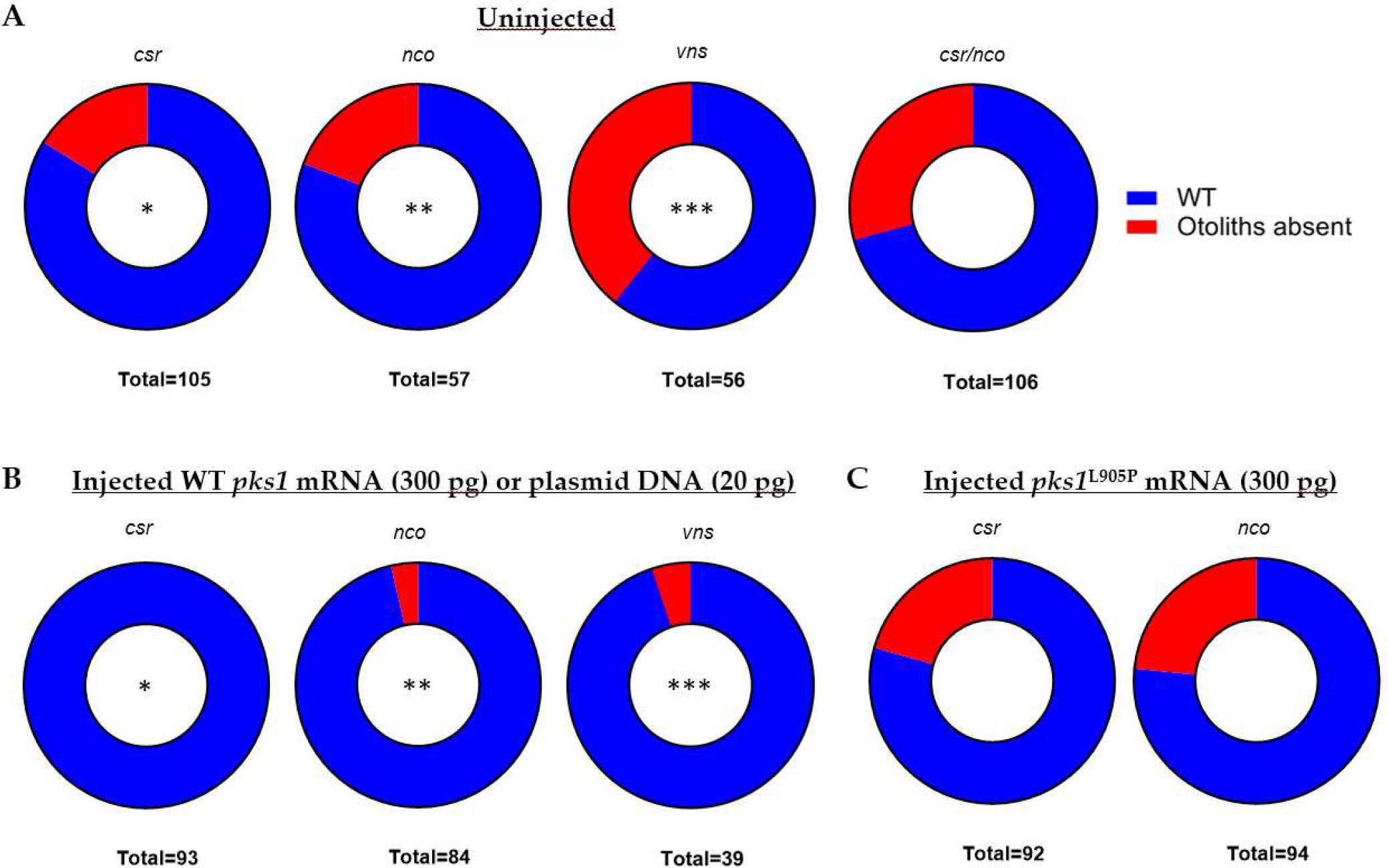
WT *pks1* nucleic acid rescues otolith formation in *csr, nco*, and *vns*. (**A**) Normal frequencies of mutant phenotypes in each uninjected strain. All four pairings follow homozygous recessive mode of inheritance. (**B**) Results of injected embryos show that Japanese medaka *pks1* mRNA (300 pg) rescues both *csr* and *nco* mutants and *pks1* DNA (20 pg) rescues *vns* mutants. (*, p < 0.0001, paired *t*-test)(**, p < 0.0032, paired *t*-test)(***, p = 0.0001, paired *t*-test),. Site-directed mutagenesis was used to introduce a conserved mutation in csr (A911P) the Japanese medaka construct (L905P) (**C**) Injection of *pks1*^L905P^ (300 pg) fails to rescue *csr* or *nco* mutant phenotypes.

### 3.5 Ingenuity pathway analysis of nco embryos

While *pks1* is thought to produce an otolith nucleation factor [21], its broader role during inner ear development is unknown. Ingenuity Pathway Analysis of nco at 24 hpf identified eNOS and Endothelin-1 signaling as the top up- and down-regulated pathways, respectively (Fig. 4A). Among the down regulated genes was *rdh12l*, a gene adjacent to *pks1*, suggesting that there is local control of transcription at that locus. *mir-92a*, the top down-regulated gene, has a predicted binding site in the 3’UTR of *rdh12l* (Fig. S2) [27]. In addition, several genes listed in the top ten up- or down-regulated lists are also enriched in adult mechanosensory hair cells such as *il11b, fosab, fosb, fosl1a, socs3a, scg5*, and *dnaaf3* (Figs. 4B-C) [28]. Of these genes, *il11b* is up-regulated during neuromast hair cell regeneration [29]. Notably, *dnaaf3* causes primary ciliary dyskinesia and morpholino knockdown of *dnaaf3* causes abnormal otolith growth [30]. While its role in inner ear development is unknown, *scg5* is expressed within the anterior and posterior poles of the otic placode during the critical period of otolith nucleation [31].

**Figure 4:**
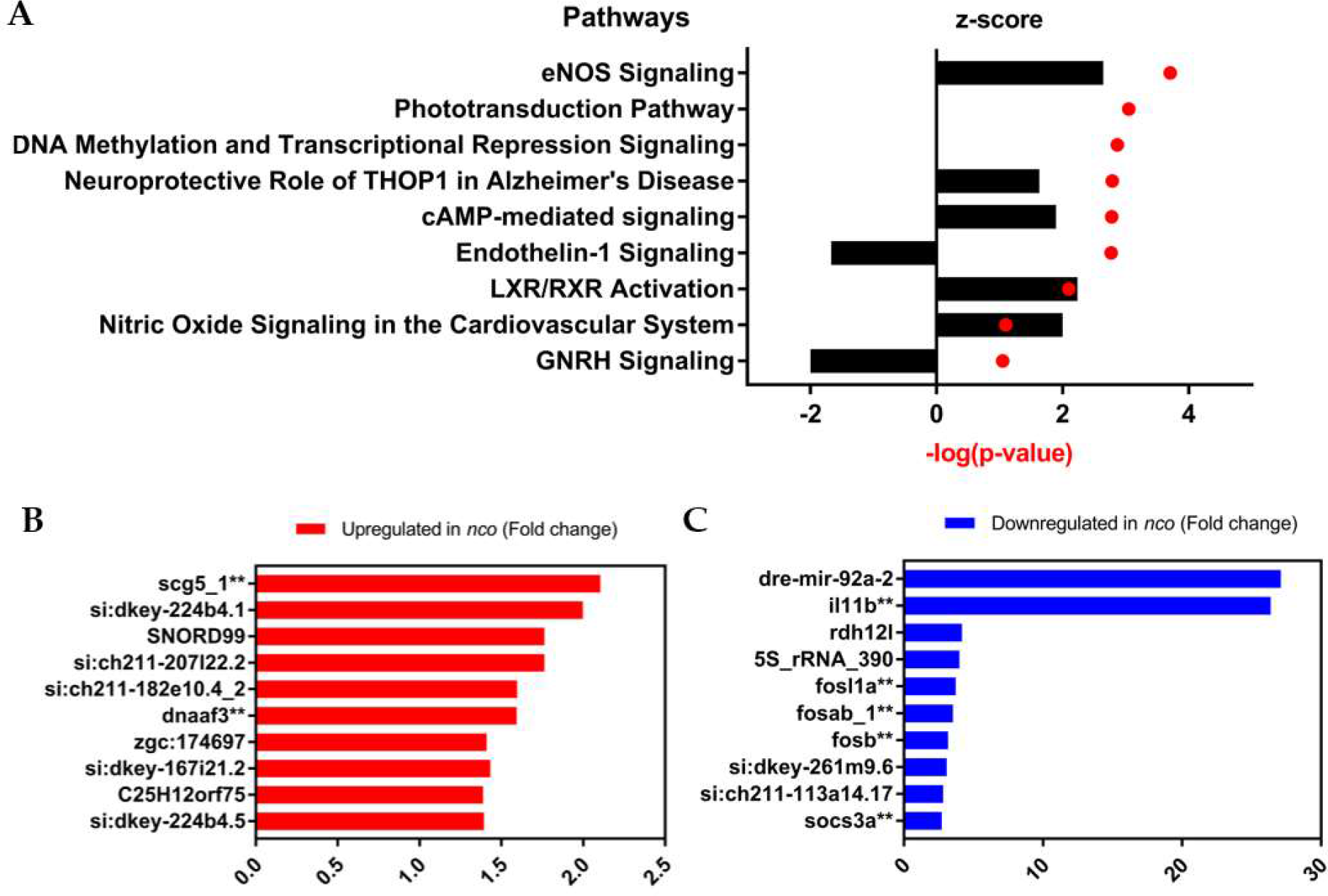
Gene expression and pathway analysis of *nco* embryos. (**A**) Ingenuity Pathway Analysis shows the top up-regulated and down-regulated pathways, which are eNOS Signaling and Endothelin-1 Signaling, respectively. Positive z-score indicated increased mRNA levels. Negative z-score indicates decreased mRNA levels. No change in mRNA levels in a z-score of zero. (**B**) Differential gene expression in the top up-regulated genes. (**C**) Differential gene expression in the top down-regulated genes. (**, expressed in adult zebrafish mechanosensory hair cells) [28].

### 3.6 Aberrant expression of proteins involved in otolith development in csr and nco

In mammalian inner ear development, Oc90 is necessary for otoconial seeding and nucleation [9, 10]. Similarly, the role of Oc90 is evolutionally conserved in zebrafish and has been previously thought to be sufficient for otolith nucleation [14]. Using immunofluorescence (IF), we saw diffuse expression of Oc90 in *csr* and *nco* otocysts (Figs. 5B-D), which demonstrated that Oc90 expression within the otocyst is not sufficient for otolith biomineralization in zebrafish. Similarly, normal localization of Otogelin (Otog), a protein required for otolith tethering in the otolithic membrane is not sufficient for Oc90 attachment. Additionally, other otoconins that are important for calcium deposition and growth were detected with diffuse expression within the otocyst such as Starmaker and Keratan Sulfate (data not shown) [32, 33].

### 3.7 Polyketide synthase expression is enriched in adult mechanosensory hair cells

Otolith nucleation is thought to be mediated by a tether-cell specific otolith precursor binding factor (OPBF), which lays the foundation for the successive biomineralization of the otolith [5, 7, 34]. The presence of an OPBF was proposed almost two decades ago and its identification proves to be elusive [34]. Recent studies suggest that one or more OPBFs are expressed by tether-cells and help to mediate otolith nucleation by binding other OPPs [5, 7, 35].

We sought to assess if *pks1* or its enzymatic product is a tether-cell OPBF. First, we demonstrated that the total number of hair cells remain unchanged during early development in *nco*, suggesting there are no differences in tether cell maturation and maintenance (Figs. 5E-G). Then, using publicly available RNA-seq data, *pks1* mRNA can be detected during the critical period of otolith nucleation [36]. Previous data has shown its localization in the otic vesicle at 19 hpf [21], supporting its role as an OPBF. *pks1* is enriched (7.46-fold increase) in mechanosensory hair cells compared to support cells within the adult zebrafish inner ear (Table S1). Additionally, RNA-seq data suggests *pks1* appears to be differentially expressed in support cells. Support cells predominantly express a 300bp region of the 5’UTR of the *pks1* transcript [28]. A search for transcriptional regulatory motifs in the 5’UTR of *pks1* found a predicted binding site for TCF-3 [37], a transcription factor highly expressed in adult mechanosensory hair cells [28]. While the role of TCF-3 in the inner ear is unknown, it is expressed within the otic vesicle during the critical period of otolith nucleation [31].

**Figure 5:**
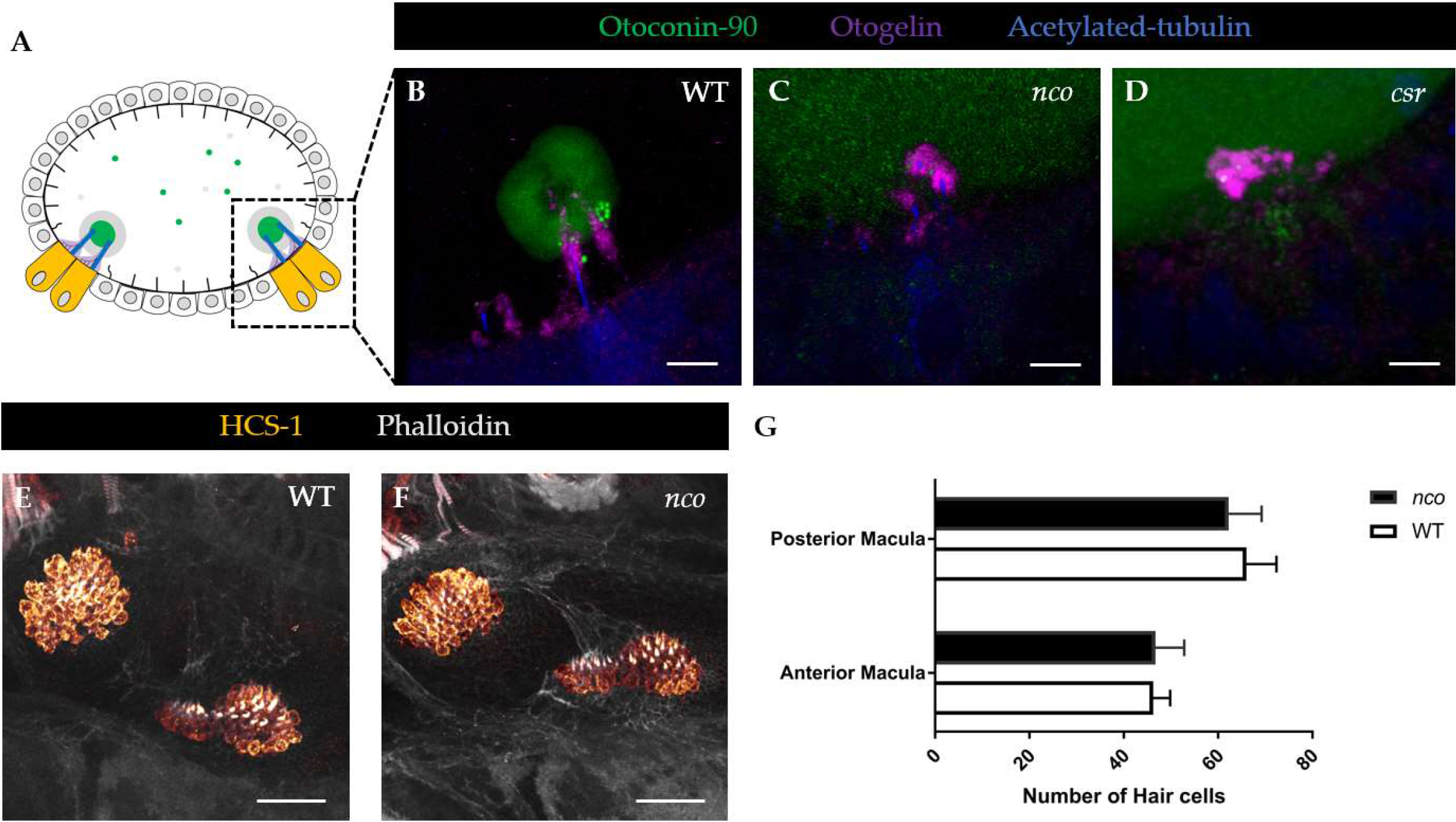
Aberrant expression of proteins invovled in otolith development in *csr* and *nco*. (**A**) Schematic of otic vesicle at 27 hpf. Anterior to right. (**B**) In WT, Oc90 is expressed the mineralized otolith, which is situated atop the otolithic membrane (Otog), at 27 hpf. Scale bar = 5 μm. (**C-D**) Oc90 has diffuse expression within the otocyst of *csr* and *nco*. *nco*, Otog is localized near the apical surface of hair cells. (**E-F**) Expression showing hair cells in WT and *nco* larvae at 5dpf. Scale bar = 25 μm. (**G**) Quantification of hair the posterior and anterior macula of WT and *nco* (n = 4).

## 4. Discussion

The mutants *csr, nco*, and *vns* were chosen for this study because each lack the necessary factors such as an OPBF for otolith seeding and biomineralization. To determine the genes responsible for otolith agenesis in these mutants, we used two complementary approaches. The first approach was Whole Genome Sequencing of the csr mutant genome to identify regions of high homology. This indeed was difficult as the csr background strain was heavily inbred, resulting in multiple peaks of high homology. Since we demonstrated *csr* and *nco* are genetically-linked, we sought to further clarify the responsible locus using a second method (i.e. RNA-seq of the *nco* transcriptome) for comparison. This result pinpointed a region of high homology near the end of the 24^th^ chromosome. While deciphering potentially deleterious mutations within that region, we focused on *pks1* following evidence that it is responsible for otolith nucleation in Japanese medaka [21]. While these species are evolutionarily divergent, the shared phenotype between medaka and our mutants suggested that the role of *pks1* is conserved. As a result, we chose to use medaka *pks1* nucleic acid to rescue otolith formation in *csr, nco* and *vns* mutants. Similarities can also be drawn with other zebrafish mutants such as *keinstein*, which has diffuse expression of Starmaker within the otocyst and exhibits similar circling swimming behaviors [38, 39]. Furthermore, *keinstein* may be another *pks1* allele due to its predicted chromosomal location [40].

While WT medaka *pks1* rescues otolith biomineralization in *csr* and *nco*, differences in penetrance of exogenous ions on otolith formation suggested the nature of each mutation is fundamentally different. This was confirmed by Sanger sequencing that *nco* has a premature stop codon while *csr* likely makes a defective protein that may be stabilized by exogenous ions. This defective protein may be the explanation for the differences in Mitotracker localization in *csr*. Due to its surface stain expression, we hypothesize that Mitotracker was localized to mitochondria-rich ionocytes [41]. Ionocytes have previously been implicated in otolith formation as mutations in *gcm2*, which is responsible for ionocyte maturation, leads to otolith agenesis [42]. We hypothesize that the endolymph in *csr* and *nco* mutants has the necessary components for otolith nucleation [2] but lack a trigger produced by *pks1*. Additionally, the absence of *pks1* does not visibly appear to affect hair cell development that are required for otolith nucleation [5]. IF of *csr* and *nco* embryos demonstrated that expression of a critical otoconial seeding protein, Oc90, within the otocyst is not sufficient for otolith biomineralization in the presence of the otolithic membrane.

One caveat is that the penetrance of otolith formation is influenced by the genetic background of zebrafish. When treated with the small molecule 31N3, WT embryos in the AB/EKW background fail to develop otoliths [6]. However, 31N3 fails to inhibit otolith formation in the TL and TU strains, suggesting that there are potential genetic modifiers that influence otolith nucleation in these backgrounds. In the case of *csr*, homozygosity at the locus is compatible with otolith agenesis in the AB background and, with proper development, in the TL background. This suggests *csr* may be a hypomorphic allele and the AB background can overcome the loss of Pks1 function with enhanced ion flux. Ironically, the mutant phenotype was lost when csr was outcrossed to the WIK background. It was only until *csr* was backcrossed to the AB background that the mutants were recovered. Altogether, we suggest that the AB background heavily influences the penetrance of otolith formation.

While *pks1* likely acts as an enzyme whose expression is enriched in adult mechanosensory hair cells [28], its product acts as an OPBF and is required for otolith nucleation in zebrafish. However, the molecular function of *pks1* remains unknown. Using *nco* RNA-seq data, we performed an Ingenuity Pathway Analysis, which identified eNOS and Endothelin-1 signaling as the most up- and down-regulated pathways, respectively. eNOS signaling could be impacted by *pks1* metabolites such as iromycin, which has been shown to inhibit this pathway [43]. Both eNOS and Endothelin-1 have been implicated in inner ear development and function. Notably, it has been demonstrated that these pathways are inversely related in sensorineural hearing loss [44]. An example of this is Waardenburg syndrome, caused by mutations in endothelins, which cause abnormal pigmentation and sensorineural hearing loss [45]. During early development, Endothelin-1 mRNA turns on during the critical period of otolith nucleation [31, 36] and is detected in the otic vesicle at 24 hpf [46]. Endothelin-1 and its receptor (*ednraa*) are both enriched in adult zebrafish inner ear support cells [28].

Additionally, Endothelin-1 has been implicated with the FOS-family of genes (*fosab, fosb, and fosl1a*) and *socs3a*, which are all differentially expressed in *nco* at 24 hpf. They are all part of a regulatory network during hypergravity-mediated bone formation [47], which might suggest a common mechanism between bone mineralization and otolith biomineralization. Future studies will attempt to clarify the roles of Endothelin-1 and eNOS signaling pathways during otolith biomineralization.

## Author Contributions

Conceptualization, K.T. and K.K.; Methodology, K.T, S.G. and K.K..; Validation, K.T., S.G., and L.H.; Formal Analysis, K.T. and S.G.; Investigation, K.T., S.G., and L.H.; Resources, J.S. and K.K; Data Curation, J.S. and K.K.; Writing-Original Draft Preparation, K.T.; Writing-Review & Editing, K.T., S.G., J.S., and K.K; Visualization, K.T. and K.K.; Supervision, J.S. and K.K.; Project Administration, J.S. and K.K.; Funding Acquisition, J.S. and K.K.

## Acknowledgments

We recognize the University of Nebraska Medical Center Genomics Core Facility for assistance with sequencing and bioinformatics. We thank Dr. Hiroyuki Takeda from the University of Tokyo for supplying the Japanese medaka *pks1* mRNA construct. We acknowledge Creighton University Integrated Biomedical Imaging Facility for assistance with confocal microscopy. Finally, we express gratitude to the members of the Kramer Lab at Creighton University for their support with zebrafish husbandry.

## Appendix A- Supplemental Material

**Figure S1:**
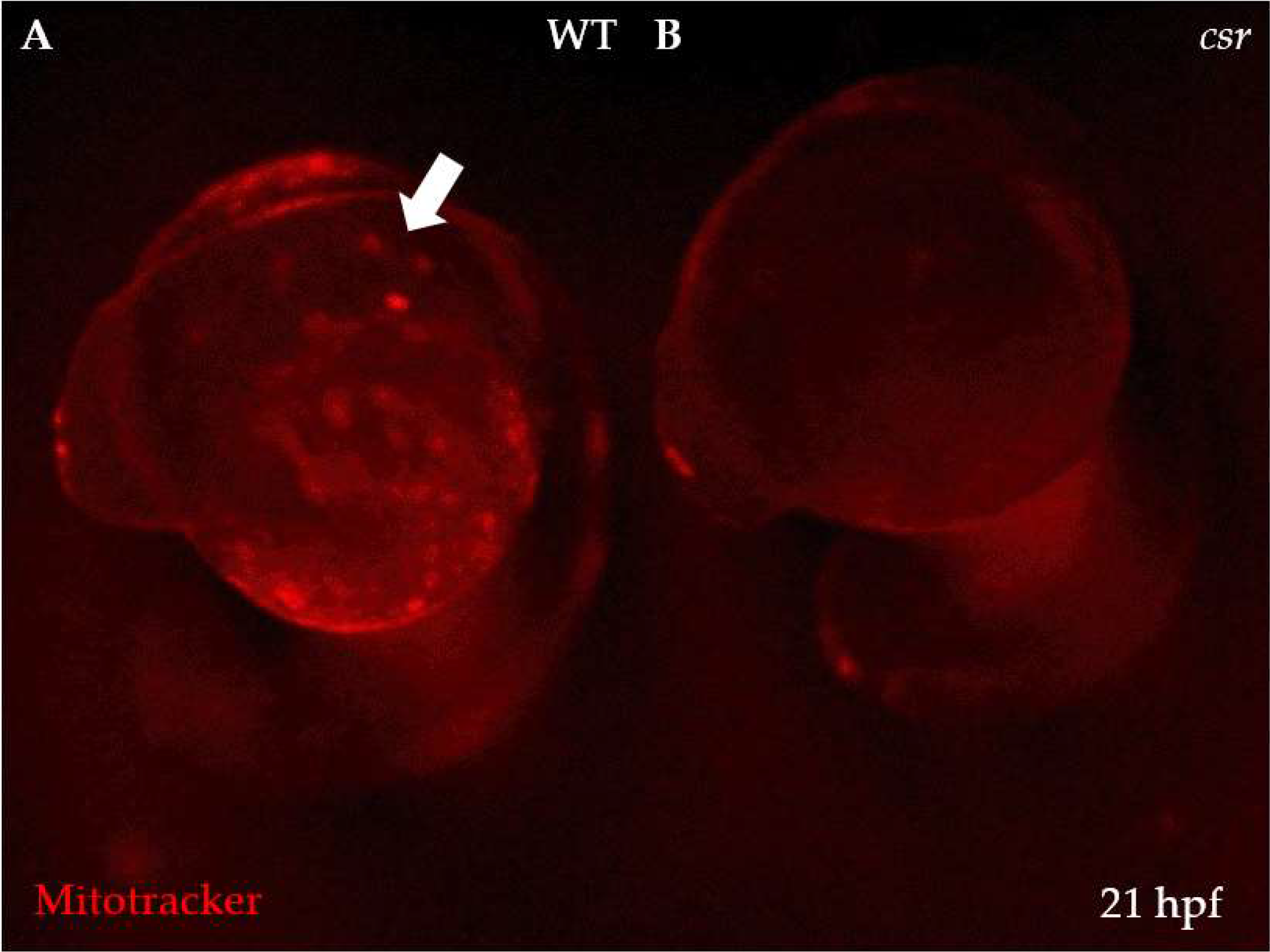
Spatial differences in mitochondrial membrane potentials. (**A**) While Mitotracker marks active mitochondria in WT, (**B**) *csr* embryos show a lack of Mitotracker expression during early development. Arrow indicates otic vesicle.

**Figure S2:**
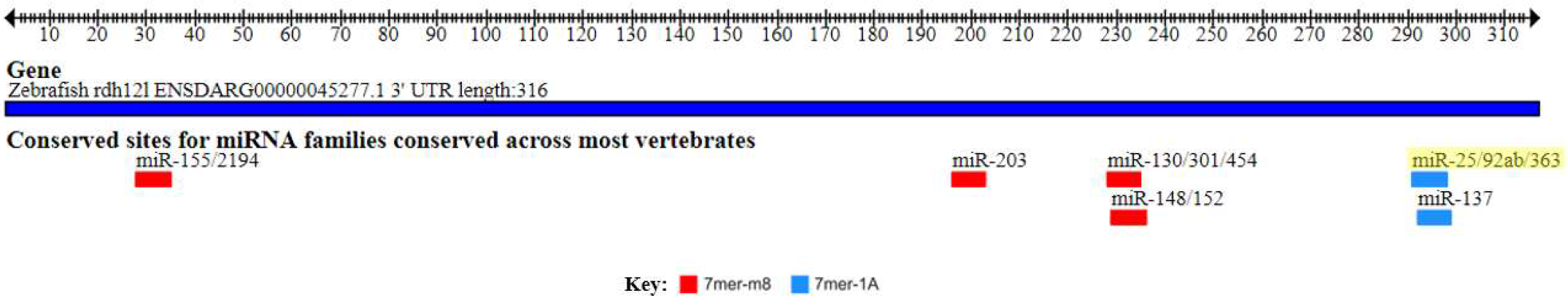
*miR-92a* binding site in the 3’ UTR of *rdh12l*. TargetScanFish 6.2 of *rhd12l* in zebrafish shows potential microRNA binding sites including *miR-92a*, which is the most down-regulated gene in *nco* embryos at 24 hpf.

**Table S1.**
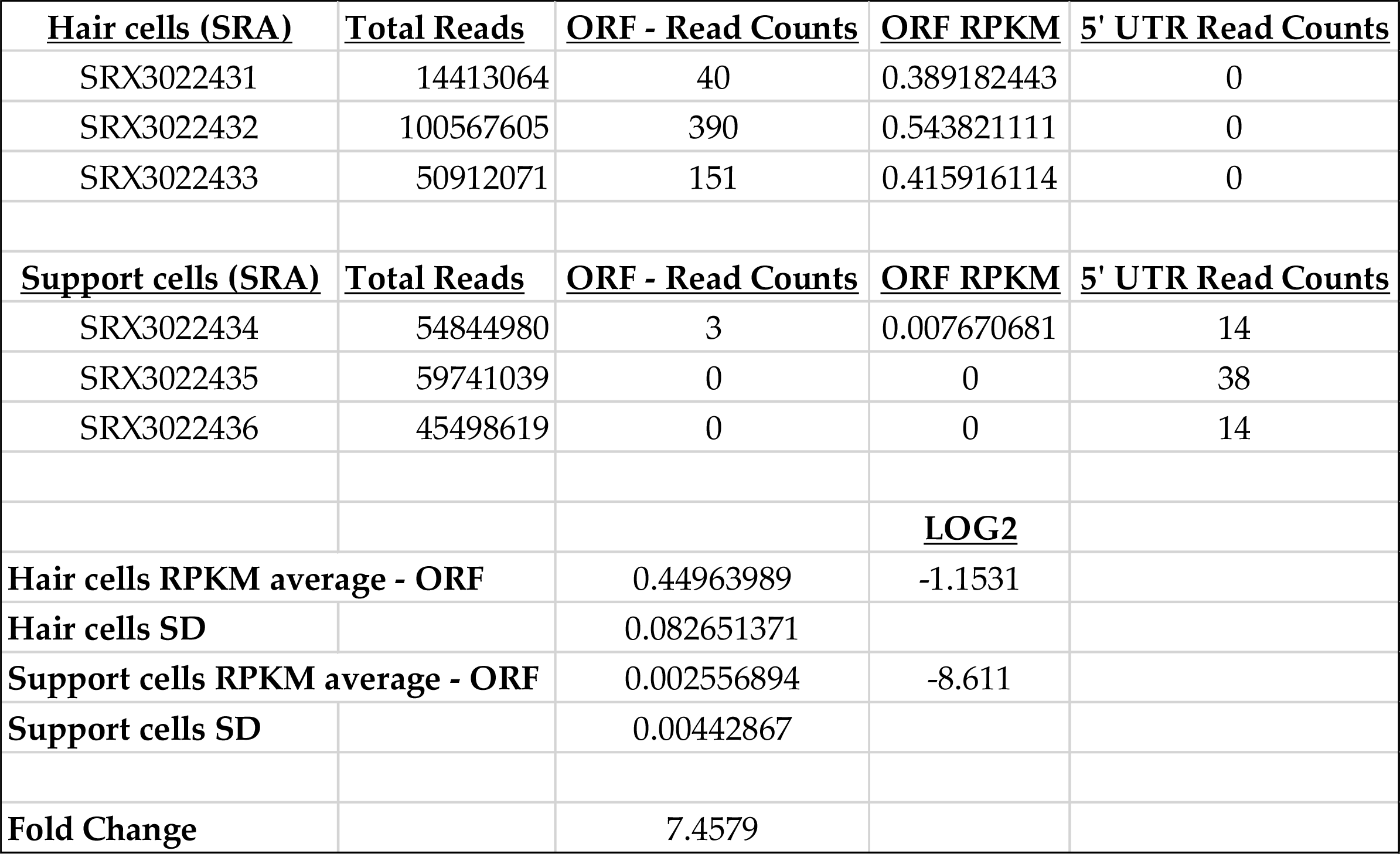
Differential expression of pks1 in hair and support cells.

| **Hair cells (SRA)** | **Total Reads** | **ORF - Read Counts** | **ORF RPKM** | **5' UTR Read Counts** |
| --- | --- | --- | --- | --- |
| SRX3022431 | 14413064 | 40 | 0.389182443 | 0 |
| SRX3022432 | 100567605 | 390 | 0.543821111 | 0 |
| SRX3022433 | 50912071 | 151 | 0.415916114 | 0 |
| **Support cells (SRA)** | **Total Reads** | **ORF - Read Counts** | **ORF RPKM** | **5' UTR Read Counts** |
| SRX3022434 | 54844980 | 3 | 0.007670681 | 14 |
| SRX3022435 | 59741039 | 0 | 0 | 38 |
| SRX3022436 | 45498619 | 0 | 0 | 14 |
|  |  |  | **LOG2** |  |
| **Hair cells RPKM average - ORF** |  | 0.44963989 | -1.1531 |  |
| **Hair cells SD** |  | 0.082651371 |  |  |
| **Support cells RPKM average - ORF** |  | 0.002556894 | -8.611 |  |
| **Support cells SD** |  | 0.00442867 |  |  |
| **Fold Change** |  | 7.4579 |  |  |

